# Microbiome-enabled genomic selection improves prediction accuracy for nitrogen-related traits in maize

**DOI:** 10.1101/2023.03.03.530932

**Authors:** Zhikai Yang, Tianjing Zhao, Hao Cheng, Jinliang Yang

## Abstract

Root-associated microbiomes in the rhizosphere (rhizobiomes) are increasingly known to play an important role in nutrient acquisition, stress tolerance, and disease resistance of plants. However, it remains largely unclear to what extent these rhizobiomes contribute to trait variation for different genotypes and if their inclusion in the genomic selection (GS) protocol can enhance prediction accuracy. To address these questions, we developed a microbiome-enabled GS (MEGS) method that incorporated host SNPs and ASVs (amplicon sequence variants) from plant rhizobiomes in a maize diversity panel under high and low nitrogen (N) field conditions. Our cross-validation results showed that the MEGS model significantly outperformed the conventional GS model for nearly all time-series traits related to plant growth and N responses, with an average relative improvement of 3.7%. The improvement was more pronounced under low N conditions (8.4% — 40.2% of relative improvement), consistent with the view that some beneficial microbes can enhance N nutrient uptake, particularly in low N fields. However, our study could not definitively rule out the possibility that the observed improvement is partially due to the ASVs being influenced by microenvironments. Using a high-dimensional mediation analysis method, our study has also identified microbial mediators that establish a link between plant genotype and phenotype. Some of the detected mediator microbes were previously reported to promote plant growth. The enhanced prediction accuracy of the MEGS models, demonstrated in a single environment, serves as a proof-of-concept for the potential application of microbiome-enabled plant breeding for sustainable agriculture.

## Introduction

An increasing number of studies have suggested that plant-associated microbial communities, especially the microbial species colonizing in the plant roots, can stimulate plant growth (Saleem et al. 2019), enhance nutrient availability in soils (Gomes et al. 2018; Zhu et al. 2016), and decrease abiotic stress responses (Hussain et al. 2018; Xu et al. 2018). Harnessing these beneficial microbes in crop production provides a promising opportunity for crop improvement to fight against climate challenges, reduce dependency on chemical fertilizers, and boost genetic gain. Indeed, from the beginning of plant domestication, rhizosphere microbes were reported to be involved in crop performance (Soldan et al. 2021). For example, studies have shown that domesticated plants exhibited distinct microbial compositions as compared to the wild ancestors and showed a reduced ability to establish symbiotic relationships with beneficial microbes (Abdelfattah et al. 2022; Abdullaeva *et al*. 2021). Recent crop improvement further reduced the microbial diversity in a number of different crop species, including wheat (Hetrick et al. 1992), maize (Sangabriel-Conde et al. 2014), and soybean (Kiers et al. 2007). Realizing the importance of microbiomes in contributing to crop production, recently, efforts have been made to screen for beneficial microbes as potential seed additives to promote plant performance (Singer et al. 2021; Yee et al. 2021). Though promising results were found in controlled environments (Eida et al. 2017; Kaur et al. 2020; Sessitsch et al. 2019), it is difficult for many microbial inoculants to survive for a long period of time under field conditions (Piromyou et al. 2011). Considering the collective genomes of microbial species colonized in plants as the secondary genome of a plant (Berendsen et al. 2012), a hologenome (i.e., plant genome with its associated endocellular or extracellular microbiome) approach to incorporate the naturally occurring genotype-specific microbiomes into the genomic selection (GS) protocol provides an alternative strategy to improve tomorrow’s crops.

GS, an innovative plant and animal breeding technology, enables the selection of promising individuals (and the associated heritable microbes) to advance to the next generation before or without phenotyping, therefore reducing the generation interval and increasing the genetic gain per unit of time. After the initial introduction of the landmark GS research conducted by Meuwissen *et al*. (2001), animal and plant breeders embraced the unprecedented acceleration of GS with the development of statistical methods and computational tools (de Koning 2016), including G-BLUP (genomic best linear unbiased prediction), RR-BLUP (ridge regression best linear unbiased prediction), and the Bayesian regression models (BayesA, BayesB, BayesC*π*, Bayes LASSO, etc.) (Krishnappa et al. 2021; Wang et al. 2015; Burgueño et al. 2012; Wang et al. 2018; Gianola et al. 2009; Habier et al. 2011). The Bayesian regression models (with different assumptions for prior distributions of marker effects) allow genome-wide markers to have different effects and variances and usually achieve high prediction accuracy (Wang et al. 2018). However, Bayesian regression models might be more sensitive to the number of QTLs as compared to the RR-BLUP and G-BLUP methods (Wang et al. 2015). In addition to these parametric methods, non-parametric methods, such as reproducing kernel Hilbert space (RKHS) (Gianola et al. 2006) and deep learning (DL) based methods (Gianola et al. 2011), are likely be more efficient for capturing non-additive genetic effects than conventional methods, while there is no clear superiority in terms of prediction power (Montesinos-López et al. 2021). Albeit the rapid development of the GS methods in the past 20 years, little attention has been paid to incorporating host-associated microbiomes into the prediction protocols (but see a recent simulation study in dairy cattle (Pérez-Enciso et al. 2021)).

For microbiome-enabled GS (MEGS) modeling, it is straightforward to consider different microbes in a linear mixed model as the random variables, similar to the SNP markers used in conventional GS. However, unlike SNPs, microbiomes are dynamic and profoundly affected by soil microenvironments and root exudates, composed of organic acids, polysaccharides, and other metabolites (Canarini et al. 2019). These differentially recruited microbiomes by different genotypes may, in turn, impact soil physicochemical characteristics and nutrient bioavailability for plants, such as nitrogen (N) availability, ultimately regulating plant physiological processes and leading to phenotypic variation. In such a scenario, microbiomes can be modeled as an intermediate process bridging the host genotype and phenotype. In human genetics studies, using Mendelian randomization analysis by treating the microbiome as an exposure, results revealed potential causal effects of the human gut microbiome on blood metabolites (Liu et al. 2022), ulcerative colitis, and rheumatoid arthritis (Kurilshikov et al. 2021), abdominal obesity (Xu et al. 2021), or even colorectal cancer (Ni et al. 2022). However, Mendelian randomization assumes that there is no other pathway through which the instrumental variables (i.e., host genotypes) affect the outcome (i.e., host phenotypes) other than the exposure itself. Violation of this assumption can introduce bias (Hemani et al. 2018). Similar to the concept of Mendelian randomization analysis but without the assumption that the effect is solely through the exposure, either directly or indirectly through the host genotypes, genome-wide mediation analysis enables the identification of significant intermediate mediators with large effects (Yang et al. 2022).

In this study, by leveraging the GS methods and our recently developed mediation models, we conducted integrative analyses using our previously published datasets (Meier et al. 2022; Rodene et al. 2022) collected on the maize diversity panel — a panel representing maize genetic diversity in temperate latitudes (Flint-Garcia et al. 2005). The rhizosphere microbiomes were collected under high N (HN) and low N (LN) field conditions (Meier et al. 2022). From the same fields, phenotypic data were collected using an unmanned aerial vehicle (UAV) in a time-series manner (Rodene et al. 2022). In our MEGS analysis, a linear mixed model was used to predict maize phenotype by including both host genotype and rhizosphere microbiome, where SNPs and ASVs were treated as random effects with different variance components. Additionally, we modeled a presumed causal chain from host genotype to host-associated microbe to host phenotype using our previously developed high-dimensional mediation analysis method (Yang et al. 2022). These methods serve as an initial trial to integrate genome and microbiome data, aiming to enhance prediction accuracy and identify beneficial microbes for use as seed additives in field conditions. Overall, our study highlights the potential of beneficial microbes in enhancing crop performance and emphasizes the importance of considering plant-microbe interactions in plant breeding.

## Materials and methods

### Microbiome and phenotype data in the maize diversity panel

The rhizosphere microbiome (or rhizobiome) data were obtained from our previously published study collected from a subset of the maize diversity panel (*n* = 230 genotypes) eight weeks after planting in both HN and LN field conditions in 2019 (Meier et al. 2022). The rhizobiome data included *n* = 3, 626 amplicon sequence variants (ASVs) that can be clustered into 154 microbial groups. These microbial groups spanned 19 major classes of rhizosphere microbiota. In summary, the relative abundances of *n* = 3, 626 ASVs were collected for *n* = 795 observations.

Meanwhile, high throughput phenotyping data were collected from the same field in a time-series manner using UAV (Rodene et al. 2022). After image analysis at the plot level, a number of vegetation indices (VIs) were obtained, some of which showed a high correlation with conventional agronomic traits, such as leaf N content and 20-kernel weight. Of the eight VIs assessed, the Visible Atmospherically Resistant Index (VARI) showed the highest overall correlations with both leaf N content and 20-kernel weight; therefore, we will use VARI as a representative trait for illustrating our theme. The environmental conditions under which VIs were collected varied, leading to differences in data quality. Here, we selected the data from 11, 21, and 35 days after rhizobiome data sampling, which shared similar high quality, for our analysis. More details of phenotypic data can be found in Rodene *et al*. (2022).

### Genotypic data processing and linkage disequilibrium (LD) pruning

The genotypic data for maize HapMap V3.2.1 (with imputation, AGPv4) were obtained from the Panzea database (https://www.panzea.org/genotypes) (Bukowski et al. 2017). Using the PLINK software (Purcell *et al*. 2007), we merged the variants on different chromosomes and retained the bi-allelic SNPs only. We then performed SNP filtration by discarding variants with the missing rate > 0.3 across lines and a minor allele frequency (MAF) < 0.05, resulting in a subset of 22.5 million SNPs. Subsequently, LD-based SNP pruning was performed by calculating linkage disequilibrium (LD) between each pair of SNPs in the window of 10 kb, and one of a pair of SNPs was removed if the LD (*R*^2^) was greater than 0.1. We then shifted the window 10 bp forward and repeated the procedure, resulting in a final subset of 770k SNPs.

### Linear mixed model for microbiome-enabled genomic prediction

We conducted the MEGS using the rrBLUP software (Endelman 2011), where *n* = 50, 000 randomly selected SNPs and *n* = 3, 626 reproducible ASVs were included simultaneously (Meier et al. 2022). Below is the model used for genomic prediction with both maize SNPs and rhizobiome ASVs:

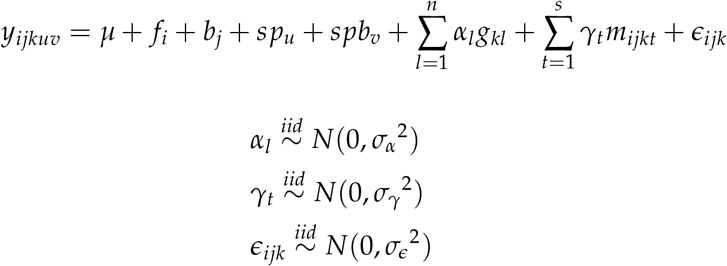

where, *y*_*ijk*_ is the observation of phenotype for the *k*th genotype in the *j*th block *u*th split plot and *v*th split plot block with the *i*th N treatment level, and there are 795 observations in total; *μ* is the intercept; *f*_*i*_ is the fixed effect of the *i*th N treatment (*i* = 1, 2); *b*_*j*_ is the fixed effect of the *j*th block (*j* = 1, 2); *sp*_*u*_ is the fixed effect of the *u*th split plot (*u* = 1, 2, 3, 4); *spb*_*v*_ is the fixed effect of the *v*th split plot block (*v* = 1, 2, 3); *α*_*l*_ is the random coefficient of the *l*th SNP; *g*_*kl*_ is the value of the *l*th SNP for *k*th genotype (*l* = 1, …, *n*, where *n* is the total number of SNPs); *γ*_*t*_ is the random coefficient of the *t*th ASV (*t* = 1, …, *s*, where *s* is the total number of ASVs); *m*_*ijkt*_ is the value (log relative abundance from 16S sequencing) of the *t*th ASV for *k*th genotype in *j*th block with the *i*th N level; and *ϵ*_*ijk*_ is the residual error.

In the model, we assumed the random coefficients of the *l*th SNP (*α*_*l*_), the *t*th ASV (*γ*_*t*_), and the residual error (*ϵ*) are independent variables following normal distributions with a mean of zero and estimated variances of 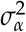 (i.e., 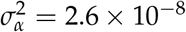 for VARI trait at 11 days after rhizobiome sampling), 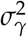(i.e., 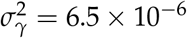 for VARI trait at 11 days after rhizobiome sampling), and 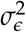, respectively. To estimate the marker effect variance 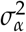, we only fitted SNPs with random effects without ASVs in an RR-BLUP model. Similarly, to get an estimate for ASV effects variance 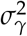, we only fitted ASVs with random effects with the first three principal components (PCs) of SNPs (fixed effects) to control for the genomic background effect in an RR-BLUP model. Finally, we assessed the predictive accuracy through randomized 5-fold cross-validation. During this process, we conducted randomization at the plot level and ensured that maize genotypes were distinct and non-overlapping between folds. And, to exclude the possibility of just adding more variables will increase prediction accuracy, we also used shuffled microbiome as control. We kept the rows of the microbiome matrix constant, with each row assigned a randomly non-repetitive sampled ID within the same N-treatment, either within or not within a quadrant.

### Conventional GS model by considering ASVs as micro-environmental factors

We conducted the following analyses to account for the microenvironmental effects on the rhizobiome. We fitted a conventional mixed linear model to obtain the BLUP value for each genotype by considering both with ASVs and without ASVs and conducted the analyses under HN and LN conditions separately.

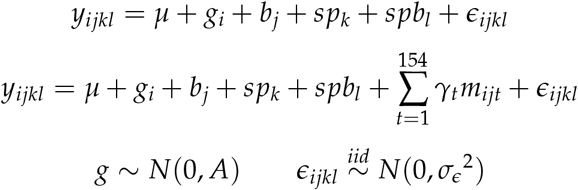

where, *y*_*ijkl*_ is the observation of phenotype for *i*th genotype in *j*th block, *k*th split plot, *l*th split plot block; *μ* is the intercept; *g*_*i*_ is the *i*th genotype effect, *i* = 1, …, 230; *A* is the additive relationship matrix calculated from SNPs using “A.mat” function from the rrBLUP package; *b*_*j*_ is the *j*th block effect, *j* = 1, 2; *sp*_*k*_ is the *k*th split plot effect, *k* = 1, 2, 3, 4; *spb*_*l*_ is the *l*th split plot block effect; *γ*_*t*_ is the random coefficient of the *t*th taxonomic group, *t* = 1, …, 154; *m*_*ijt*_ is the value of the *t*th taxonomic group for *i*th genotype in *j*th block; *t* = 1, …, 154; and *ϵ*_*ijkl*_ is the residual error. We employed the randomized 5-fold cross-validation to obtain the prediction accuracy.

### Mediation analysis by considering microbes as intermediate variables

Mediation analysis introduces a type of variable called mediator to infer the underlying mechanism of the relationship between an independent variable and a dependent variable (Baron and Kenny 1986). Similar to our previous study (Yang et al. 2022; Zhang 2021), here we conducted genome-wide mediation analysis using SNPs as exposures, ASVs as microbe mediators, and the first three principal components (PCs) with other experiment and treatment factors as confounders. The mediation analysis consists of two models: the mediator model and the outcome model. Specifically, the mediator model is shown as follows:

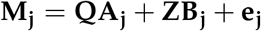

where, **M**_**j**_ (a *n ×* 1 vector) represents the abundance of the *j*th ASV; **Q** (a *n×* (*s* + *f* + *b* + *u* + *v*) matrix) is the design matrix for first *s* PCs of genotypes, *f* nitrogen treatments *b* blocks, *u* split plots and *v* split plot blocks (*s* = 3, *f* = 2, *b* = 2, *u* = 4, *v* = 3 in our analysis); **A**_**j**_ (a (*s* + *f* + *b* + *u* + *v*) *×* 1 vector) is the coefficients of the first *s* PCs, *f* nitrogen treatments, *b* blocks, *u* split plots and *v* split plot blocks to the *j*th ASV; **Z** (a *n ×* matrix) represents the SNP set of the population composed of *n* individuals with *q* number of SNPs; **B**_**j**_ (a *q ×* 1 vector) is the coefficients of the *q* SNPs to the *j*th ASV (*j* = 1, …, *p*); and **e**_**j**_ is the vector of the residual errors with 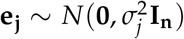.

Additionally, we fitted the SNPs and the microbe mediators (i.e., ASVs) in the outcome model, which uses phenotype as the response variable, as shown below:

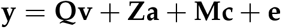

where, **y** (a *n ×* 1 vector) represents the phenotype; **Q** (a *n ×* (*s* + *f* + *b* + *u* + *v*) matrix) and **Z** (a *n × q* matrix) are the same matrices as the above mediator model; **v** (a (*s* + *f* + *b* + *u* + *v*) *×* 1 vector) is the coefficients of the first *s* PCs, *f* nitrogen treatments, *b* blocks, *u* split plots and *v* split plot blocks; **a** (a *q ×* 1 vector) denotes the coefficients of the SNPs to the phenotype; **M** (a *n × p* matrix) is the abundance of the ASVs (i.e., a matrix combines all the vectors of **M**_**j**_); **c** (a *p ×* 1 vector) is the coefficients of ASVs to the phenotype; **e** is the vector the residual errors with **e** ∼ *N*(**0**, *σ*^2^**I**_**n**_).

## Results

### Rhizobiome incorporation into GS model improves prediction accuracy for host phenotype

Tens of thousands of microbial species colonize plant roots, many of which are heritable (Peiffer et al. 2013; Meier et al. 2021). To assess if rhizosphere microbiomes (hereafter referred to as rhizobiomes) could be leveraged to predict the plant phenotypic performance, we analyzed the composition of rhizobiomes quantified in a diversity panel of 230 maize inbred lines under high N (HN) and low N (LN) field conditions (Meier *et al*. 2022). Additionally, we obtained the image-extracted vegetation index traits collected from the same field in 2019 (Rodene et al. 2022). Since most of the vegetation index traits exhibit high correlation (see **Figure S1**), we focused on one of the most representative vegetation indices — Visible Atmospherically Resistant Index (VARI) — collected on three dates, i.e., 11, 21, and 35 days after microbiome sampling (DAMS) in the field.

We fitted the host rhizobiomes (ASVs) at varying levels of resolution — 3,626 ASVs, 154 taxonomic groups, and 19 classes — along with host genotypes (SNPs) as explanatory variables, by incorporating them as random effects into a linear mixed model (**Materials and Methods**). After conducting 20 randomized five-fold crossvalidations, our results indicate that including ASVs in the model significantly enhances the GS prediction accuracy compared to the traditional SNP-only model for all three time points when using either 3,626 ASVs (**Figure 1B**) or 154 taxonomic groups (**Figure 1B**). For the 19 classes (**Figure 1C**), a slight increase in prediction accuracy was observed on Day 11 post-microbiome sampling, which was an anticipated result given the substantial decrease in microbiome resolution from 3,626 ASVs to 19 classes. Interestingly, the 154 taxonomic groups are as informative as the 3,626 ASVs, particularly for the 35 DAMS.

**Figure 1.**
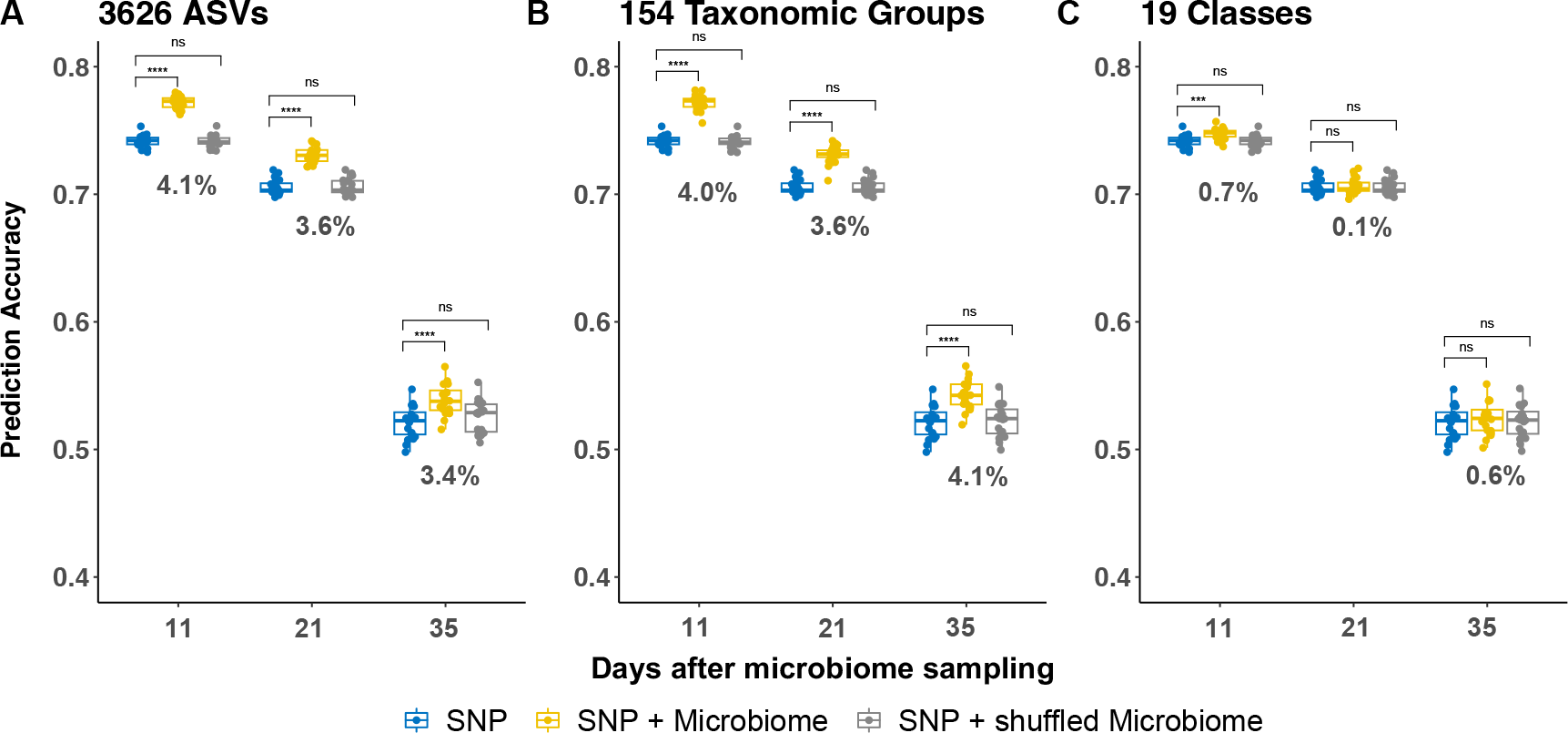
Comparison of prediction accuracy for an image-extracted trait using different variables in GS models across gradient rhizobiome resolutions. Genomic prediction results using SNPs only (blue), both SNPs and rhizobiomes (yellow), and both SNPs and shuffled rhizobiomes (grey) at different resolutions: 3,626 ASVs (**A**), 154 taxonomic groups (**B**), and 19 classes (**C**). Asterisks indicate the statistical significance of the difference in accuracy between the models: ns (not significant), * (0.01 < p-value ≤0.05), ** (0.001 < pvalue ≤0.01), ***(0.0001 < p-value ≤0.001), and **** (p-value ≤0.0001). The numbers below the boxplots are the improvement of prediction accuracy (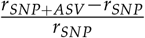, where *r*_*SNP*+*ASV*_ is the prediction accuracy for the SNP and ASV model and *r*_*SNP*_ is the accuracy for the SNP-only model) on different dates.

As the days post-microbiome sampling increased, we noted a decline in overall prediction accuracy, likely due to the decreased heritability of VARI traits as plants close to senescence. To verify that the improved prediction accuracy was not merely a consequence of a larger number of explanatory variables, we generated a comparable set of dummy variables by randomly shuffling the ASVs using different shuffling strategies (**Materials and Methods**). Results showed that the additional shuffled variables either did not significantly influence the prediction accuracy (**Figure 1**) or improved prediction accuracy but less than the original variables (**Figure S2**, this shuffling strategy within the quadrant is closer to original order). Compared to the conventional GS model utilizing only SNPs, the addition of ASVs resulted in accuracy improvements of 4.1%, 3.6%, and 3.4% at 11, 21, and 35 DAMS, respectively (**Figure 1A**). Even when measured against results from the randomly shuffled ASVs, an average improvement of 3.7% was observed. Using 154 taxonomic groups, the resulting improvement is similar or even slightly higher, with an average improvement of 3.9% (**Figure 1B**). Similar results were also observed for other image-extracted traits (see **Figure S3**).

### Prediction improvement is predominantly observed under LN conditions

To investigate the impact of N treatment on MEGS results, we fitted the ASV-based model separately for HN and LN conditions. Under HN conditions, the model did not outperform the conventional SNP-only model (**Figure 2A**). However, under LN conditions, significant improvements were observed on all three days (**Figure 2B**). Specifically, prediction accuracy increased from 55.9% to 59.7% at 11 DAMS (6.8% improvement), from 52.3% to 56.7% at 21 DAMS (8.4% improvement), and from 24.1% to 33.8% at 35 DAMS (40.2% improvement). While the rhizobiome is likely influenced by microenvironment, to account for its effects, we fitted ASVs as environmental factors in a conventional GS model to obtain the BLUP values (**Materials and Methods**). As shown in **Figure S4**, considering ASV in BLUP calculation yielded consistent results with the MEGS model (except for the 11 DAMS under HN conditions), and the results were more substantial and statistically significant under LN conditions. These findings suggest that rhizobiomes are not entirely confounded by microenvironments, as evidenced by the consistent experimental layout under both N conditions. To further test rhizobiome’s effect on host phenotype prediction, we conducted a parallel analysis using four yield component traits (cob length, cob weight, cob width, and 20-kernel weight) collected from the same field (Palali Delen et al. 2023). MEGS results revealed significant improvements for two yield-related traits (cob length and cob weight) under HN and three traits (cob length, cob weight, and cob width) under LN conditions (**Figure S5**). Similar to VI traits, the enhancements for cob length and cob weight were more pronounced under LN than HN conditions (**Figure S5**).

**Figure 2.**
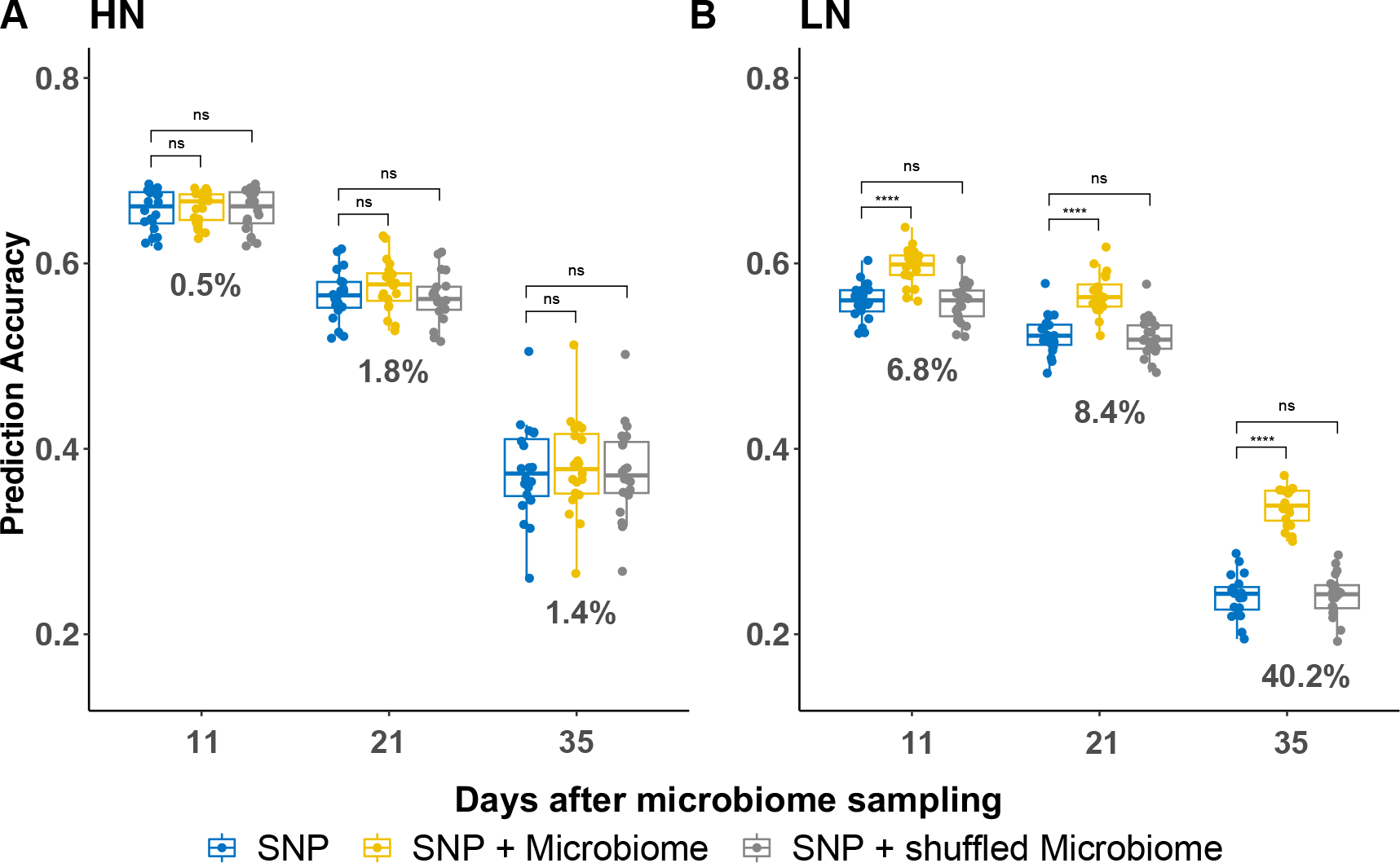
Prediction accuracy of incorporating rhizobiome under different N conditions. The prediction accuracy of using SNPs only (blue), using both SNPs and rhizobiomes (yellow), and using both SNPs and randomly shuffled rhizobiomes (grey) in high N (**A**) and LN (**B**) conditions. Asterisks indicate the statistical significance of the difference in accuracy between the models: ns (not significant), * (0.01 < p-value ≤ 0.05), ** (0.001 < p-value ≤ 0.01), ***(0.0001 < p-value ≤ 0.001), and **** (p-value ≤ 0.0001).

### Microbiomes exhibit more pronounced effects in the early days after microbiome sampling

To better understand the impact of specific microbes on prediction, we evaluated the effect sizes of the ASVs and SNPs included in the prediction model (**Figure 3**). As expected, the overall SNP effects remained largely consistent across different dates (**Figure 3B**). In contrast, the effects of ASVs diminished with the increasing interval between the initial microbiome sampling and the subsequent collection of plant phenotypes (**Figure 3A**). The investigation into the relationship between the effect sizes of ASVs and their respective taxonomic groups revealed highly significant results (Kruskal-Wallis test, p-value < 2.2 *×* 10^−16^) for both N conditions on all three days, suggesting a disproportionate enrichment of large effect ASVs within certain taxonomic groups. Notably, among the top 1% large effect ASVs (**Supplementary Table S1**), four ASVs from the taxonomic groups *Massilia niabensis, Microbacterium testaceum, Pseudomonas parafulva*, and *Sphingomonas limnosediminicola* consistently exhibited relatively large effects on VARI trait across all dates in both N conditions. Additionally, we identified one ASV belonging to *Pseudomonas parafulva* that showed a consistently large effect under HN conditions and another ASV associated with *Solirubrobacter* that consistently showed a large effect under LN conditions.

**Figure 3.**
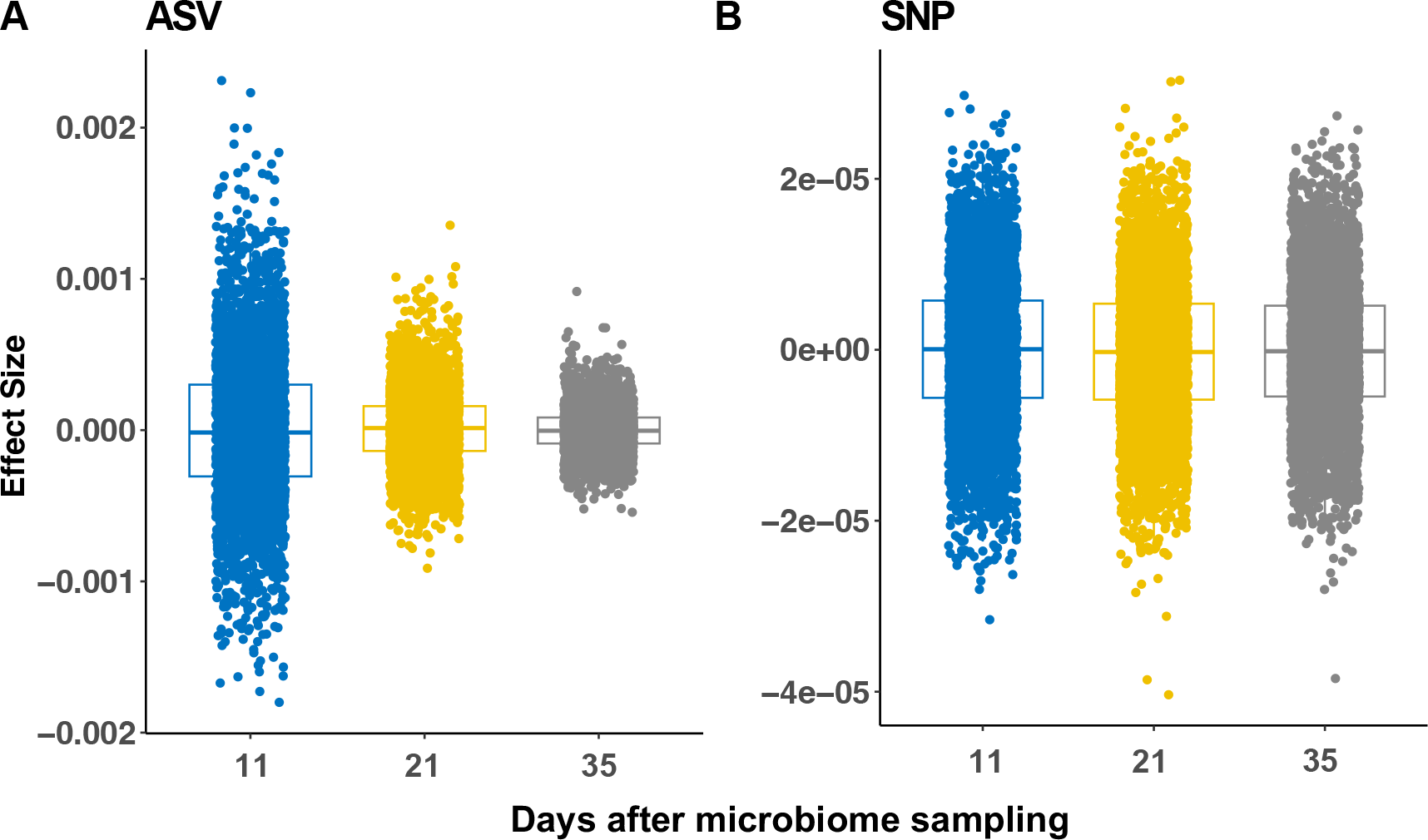
The effect sizes of ASVs and SNPs on the VARI phenotype. The distribution of effect size of ASVs (**A**) and SNP (**B**) at different DAMS.

To identify shared features of ASVs with the largest effects, we compared the top 1% of ASVs against all other ASVs in terms of their heritability (the degree to which that ASV abundance is determined by host genetics as compared to the surrounding environmental factors) and selection scores (the magnitude of effect of ASV to the phenotypic trait of interest), as calculated previously (Meier et al. 2022). The results showed that the top 1% ASVs were more heritable under LN field conditions (Wilcoxon rank sum test, p-values = 1.2 *×* 10^−4^, 0.0015, and 0.0071 for Day 11, 21, and 35, respectively under LN; p-values = 0.99, 0.15, and 0.14 for Day 11, 21, and 35, respectively under HN), and they had significantly higher selection scores under LN conditions on 11 and 21 DAMS (Wilcoxon rank sum test, p-values = 0.0019, 3.5 *×* 10^−6^, and 0.291 for Day 11, 21, and 35, respectively under LN; p-values = 0.90, 0.12, and 0.62 for Day 11, 21, and 35, respectively under HN) (**Figure S6**). These findings align with the observed enhancement in prediction accuracy in the LN field attributed to the rhizobiome and imply that plant hosts may possess a genetic mechanism that promotes the recruitment of specific microbes to alleviate LN stress.

### Microbiome mediation analysis reveals promising microbial mediators

In order to establish a causal chain from plant genotype to microbiome to plant phenotype, we sought to model ASVs as the intermediate mediators that are selectively recruited by different plant genotypes and have a significant effect on plant phenotype. Therefore, we supplemented the MEGS with our previously developed genome-wide mediation analysis (Yang et al. 2022). This approach (see **Materials and Methods**) enabled us to identify 17 unique ASVs acting as mediators for eight VI traits (**Supplementary Table S2**), with 8/17 ASVs mediating the VARI trait. Notably, ASVs in taxonomy groups of *Enterobacter, Massilia putida, Pseudomonas parafulva, TM7a sp 2* were also the top 1% with the largest effect sizes in the MEGS model under both N conditions. Additionally, the same ASVs annotated as *Massilia putida* and *Pseudomonas parafulva* were also in the top 1% large effect ASVs.

To investigate the effects of *Massilia putida* and *Pseudomonas parafulva* on plant phenotype, we performed Spearman’s rank correlation test to determine if there was a significant correlation between the abundance of ASVs and the VARI phenotype. For *Massilia putida* (**Figure 4A**), under HN conditions, we found barely negative correlations on 11 (*r* = −0.10, p-value = 0.044), 21 (*r* = −0.048, p-value = 0.34), and 35 (*r* = −0.014, p-value = 0.78) days. Under LN conditions, results showed significantly larger and positive correlations on days 11 (*r* = 0.17, p-value = 9.7 *×* 10^−4^) and 21 (*r* = 0.17, p-value = 3.4 *×* 10^−4^), yet the correlation on 35

**Figure 4.**
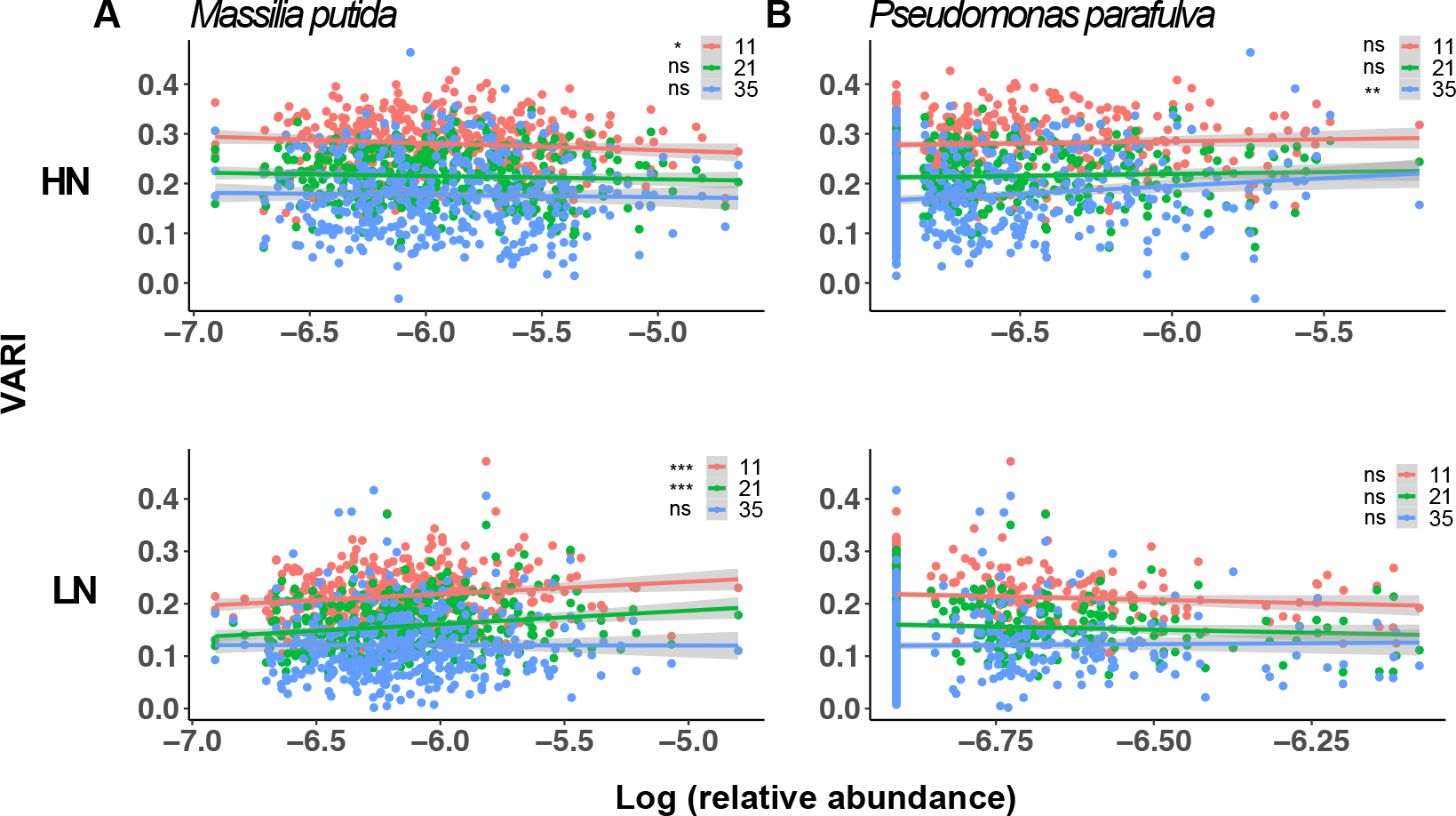
The relative abundance of *Massilia putida* and *Pseudomonas parafulva* and their relationships with the VARI phenotype. The Spearman’s rank correlations between the relative abundance of *Massilia putida* (**A**) and the VARI phenotype collected on 11, 21, and 35 DAMS under HN and LN conditions. The Spearman’s rank correlations between the relative abundance of *Pseudomonas parafulva* (**B**) and the VARI phenotype collected on 11, 21, and 35 DAMS under HN and LN conditions. Asterisks indicate the statistical significance of the difference in accuracy between the models: ns (not significant), * (0.01 < p-value ≤ 0.05), ** (0.001 < p-value ≤ 0.01), ***(0.0001 < p-value ≤ 0.001), and **** (p-value ≤ 0.0001).

DAMS was insignificant (*r* = 0.015, p-value = 0.78). These results are consistent with the previous finding that *Massilia putida* promotes plant growth under LN conditions (Yu et al. 2021). In terms of *Pseudomonas parafulva* (**Figure 4B**), an opposite trend was observed. There were larger and more significant correlations under HN conditions: Day 11 (*r* = 0.079, p-value = 0.11), 21 (*r* = 0.058, p-value = 0.25), and 35 (*r* = 0.13, p-value = 0.0083) than LN conditions: Day 11 (*r* = −0.063, p-value = 0.21), 21 (*r* = −0.046, p-value = 0.37), and 35 (*r* = 0.054, p-value = 0.29), aligning with the fact that *Pseudomonas parafulva* was top 1% ASVs under HN conditions.

## Discussion

In this study, we developed a microbiome-enabled genomic selection (MEGS) method and provided empirical evidence, albeit from only one location, that incorporating microbiome data led to a significant increase (about 4%) in prediction accuracy for most traits extracted from UAV images across different dates. We acknowledged that there might be correlations between SNPs (host genome) and ASVs (microbiome) or microenvironments and ASVs, leading to multicollinearity, but this won’t affect the screen for beneficial ASVs as seed additives for improving crop production. Besides, the spatial and temporal stability of the microbiome affects the efficacy of prediction models (Pérez-Enciso et al. 2021). Several studies suggest that the microbiome residing in the roots undergo alterations throughout the lifespan of an individual plant (Mougel et al. 2006; Houlden *et al*. 2008; Yu et al. 2012; Chaparro et al. 2014). However, research has also indicated that the microbiome could potentially reach a relatively stable condition after the initial two weeks of growth (Ibekwe and Grieve 2004; Edwards et al. 2015; Aleklett et al. 2022). Consequently, to enhance the accuracy of predictions, incorporating microbial data collected during a pre-determined stable stage would be recommended. This preference for stable stage significantly restricts the application of microbiome in prediction within breeding programs when compared to genomic data, which remains unchanged (Pérez-Enciso et al. 2021). Nonetheless, significant potential applications remain, given the observed increase in prediction accuracy (about 4% on average) after incorporating microbiome data into prediction models. This improvement is noteworthy when compared to the approximately 1% yearly genetic gain achieved using traditional breeding approaches. As the cost of obtaining ASV data through 16S rRNA sequencing decreases with advances in sequencing technology, MEGS offers an unprecedented opportunity to predict complex traits, such as N or water usage efficiency, by including ASVs as additional explanatory variables.

We analyzed prediction accuracy separately under HN and LN conditions and found that microbiome data were more beneficial under the latter (improvement can be up to 40% under N deficit field conditions). This observation is consistent with the idea that the symbiotic relationship between plants and their associated microbiomes has evolved over a long history of coexistence. However, recent changes in farming practices, especially modern crop production under N-sufficient field conditions with inorganic N fertilizer, may have a homogenizing effect on the root-associated microbiome and reduce plants’ reliance on rhizosphere microbial species for assistance in N uptake. This change is reflected in our results, where microbiome data were not as strong predictors under HN conditions compared to LN conditions. It suggests that plants may selectively recruit specific microbes to enhance nutrient absorption when soil N levels are insufficient. However, bulk soil data was not collected; therefore, a comparison of the homogeneity between the HN and LN fields cannot be conducted in this study.

Our mediation analysis, which considered the microbes as intermediaries between plant genotype and plant phenotype, pinpointed several microbe mediators. Among these, two mediator microbes, namely *Massilia putida* and *Pseudomonas parafulva*, played crucial roles in predicting the plant phenotype in the MEGS analysis. Notably, *Massila putida* was enriched in the rhizosphere and is thought to enhance plant growth and N uptake by inducing lateral root formation under LN conditions (Yu et al. 2021). And, *Pseudomonas parafulva* have previously been reported to promote plant growth (Oteino et al. 2015; Preston 2004). We also detected several other large-effect ASVs, such as *Bacillus fumarioli*, some of which have been linked to plant development (Kumar et al. 2012). Other top 1% large effect ASVs in the prediction model, like strains belonging to *Bacillus* which were only detected on certain dates, may also be important candidates for seed additives. However, further phenotypic and functional validation of these identified microbe mediators and large-effect microbes is necessary to reveal their effects on the host plants.

Due to computational constraints, we randomly sampled a subset of SNPs in low LD across the whole genome to fit the MEGS models. To determine the influence of SNP size on prediction accuracy, we performed a sensitivity test and found no significant difference in prediction accuracy when random sampling 10k, 25k, and 50k SNPs out of the 770k low LD SNPs. The prediction accuracy was reasonably high, ranging from 52.1%-76.4% for both SNP-only and SNP-ASV models across different dates (**Figure S7**). To maintain high fidelity under the current computation resource, we chose 50k SNPs for all traits and dates in the study. However, the use of more computationally efficient methods is necessary to handle larger SNP datasets in the future. Although the present study was able to achieve reasonable prediction accuracy with a limited set of SNPs and the whole set of ASVs, the use of more comprehensive datasets with more host diversity, more environmental conditions, and enhanced microbiome sampling would undoubtedly improve the accuracy of the MEGS models.

There are several caveats to microbiome-enabled prediction. One limitation is the transferability of models, as microbiomes collected in one environment at a particular plant developmental stage may differ significantly from other microbiome datasets. In fact, our previous study has demonstrated that collection date, crop rotation, and N treatment all exert significant effects on microbiome composition (Meier et al. 2021). In the current dataset, our microbiome data were collected within a three-day window at a single location. In the analysis, we controlled for microenvironment effects using the experimental design factors and also compared models with and without including ASV in calculating BLUP values. From the results, we could not determine whether the rhizobiome is simply associated with the microenvironment or actively recruited by the host genotype. Nevertheless, under both scenarios, incorporating root-associated microbiome data consistently improved prediction accuracy, especially under LN field conditions. Another foreseeable caveat is the method implementation. One of the most significant advantages of GS is the ability to predict phenotypes without planting, but microbiome data have to be collected from plant roots, which limits the extent to which predictions can be made before observations. While it may not scale up the breeding process as effectively as SNP-based GS, it still offers valuable applications, i.e., making crossing decisions before observing the phenotypes. Despite these limitations, the proof-of-concept of this study positions MEGS as a possible alternative for sustainable crop improvement. Our results suggest that microbial effects are substantial when considered close to the date of collection, but these effects diminish over time. To successfully implement this approach in a breeding program, meticulously designed experiments are essential for in-field microbiome data collection. This is crucial to mitigate microenvironmental effects and ensure uniform data collection within a specified timeframe. In addition to root-associated microbiome data, collecting the microbiome in bulk soil, along with other environmental factors such as soil type, temperature, and precipitation, is important.

## Data and Code Availability

The data and code used for the analyses can be accessed through GitHub (https://github.com/ZhikaiYang/GP_microbiome).

## Acknowledgements

This work is supported by the Agriculture and Food Research Initiative Grant number 2019-67013-29167 and 2022-67013-3656 from the USDA National Institute of Food and Agriculture, and by the U.S. Department of Energy under Award Number DE-SC0023138. This work is conducted using the Holland Computing Center of the University of Nebraska-Lincoln Start-up, which receives support from the Nebraska Research Initiative. We appreciate the constructive suggestions from anonymous reviewers.

## Supplemental Tables

**Table S1** Top 1% ASVs detected by the MEGS model. (https://github.com/ZhikaiYang/GP_microbiome/blob/master/data/supplementary/Supplemental_Table_S1_top_one_percent_asvs.txt)

**Table S2** Mediators ASVs identified by mediation analysis. (https://github.com/ZhikaiYang/GP_microbiome/blob/master/data/supplementary/Supplemental_Table_S2_mediator_asvs.txt)

## Supplemental Figures

**Figure S1.**
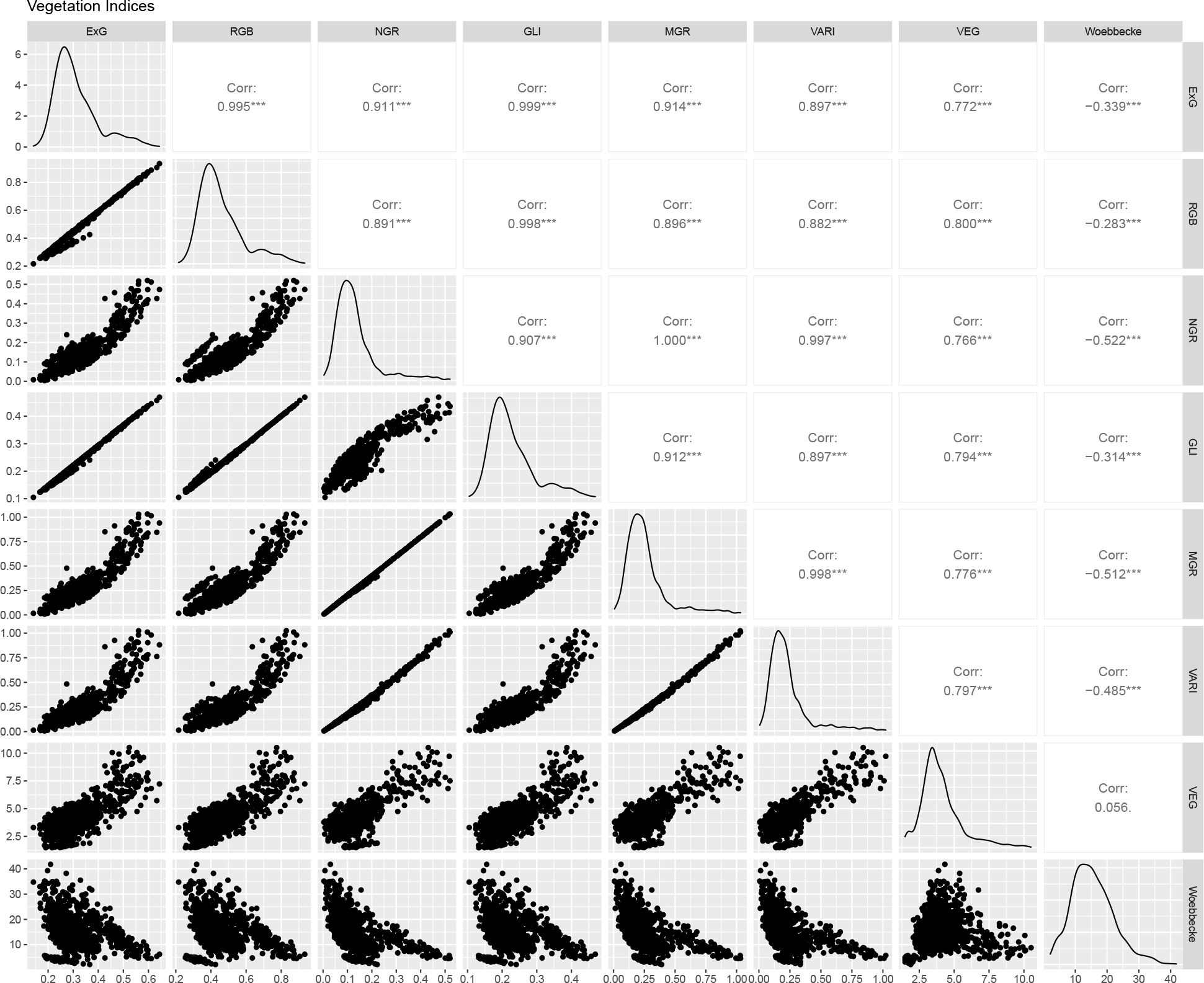
The distributions and correlations of the eight vegetation indices (VI). The upper right panel shows the pair-wise correlations, the lower left panel shows the corresponding scatter plots of each VI, and the diagonal shows the density plots of each VI. The statistical significance of the Pearson correlation is indicated by asterisks: * (0.01 < p-value ≤ 0.05), ** (0.001 < p-value ≤ 0.01), ***(0.0001 < p-value ≤ 0.001), and **** (p-value ≤ 0.0001).

**Figure S2.**
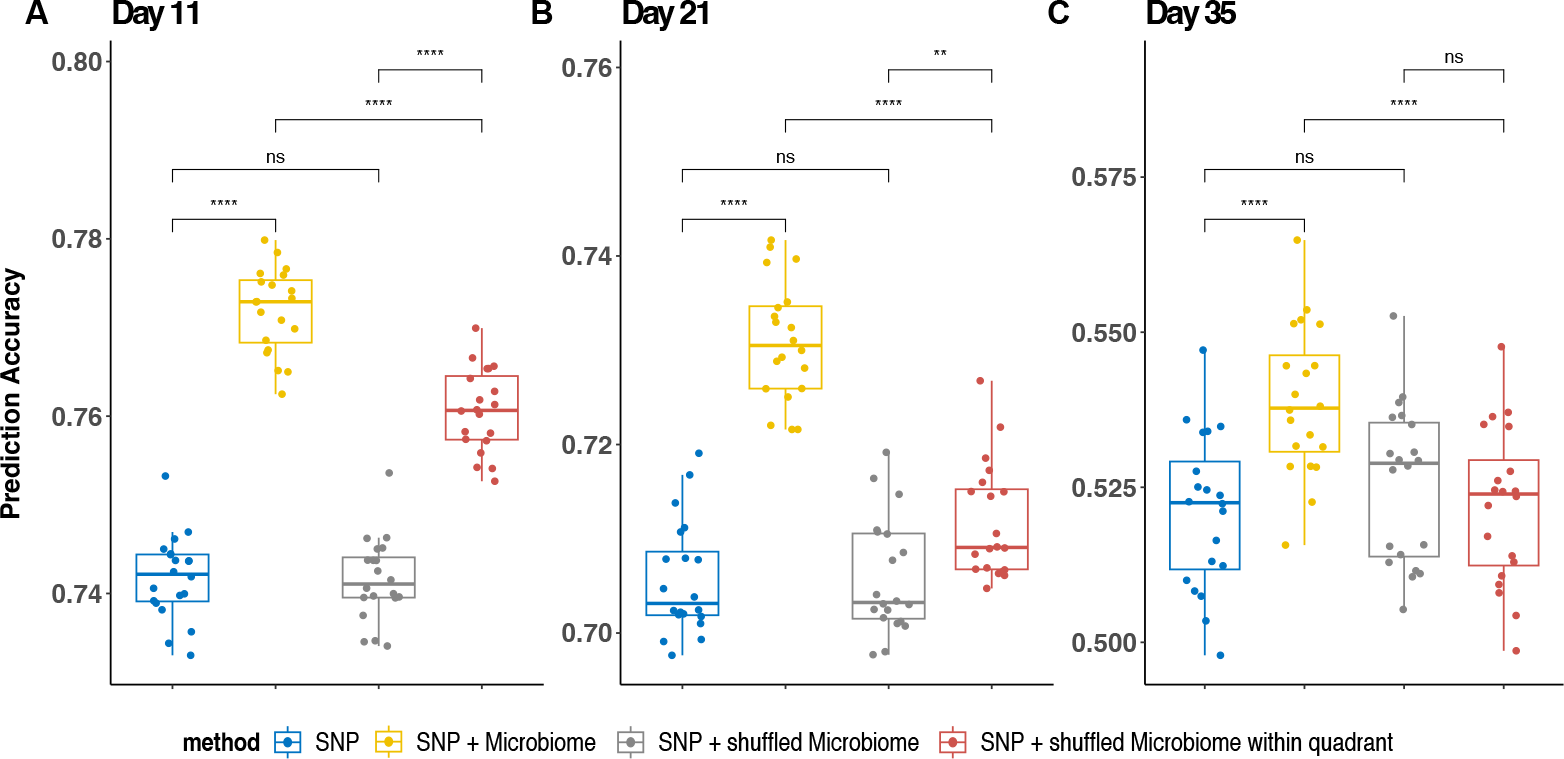
Comparison of prediction accuracy using different shuffling strategies. Genomic prediction results using SNPs only (blue), both SNPs and microbiome (yellow), both SNPs and shuffled microbiome (grey, shuffling within N-treatment but not within quadrant), and both SNPs and shuffled microbiome within quadrant (red). Asterisks indicate the statistical significance of the difference in accuracy between the models: ns (not significant), * (0.01 < p-value ≤ 0.05), ** (0.001 < p-value ≤ 0.01), ***(0.0001 < p-value ≤ 0.001), and **** (p-value ≤ 0.0001).

**Figure S3.**
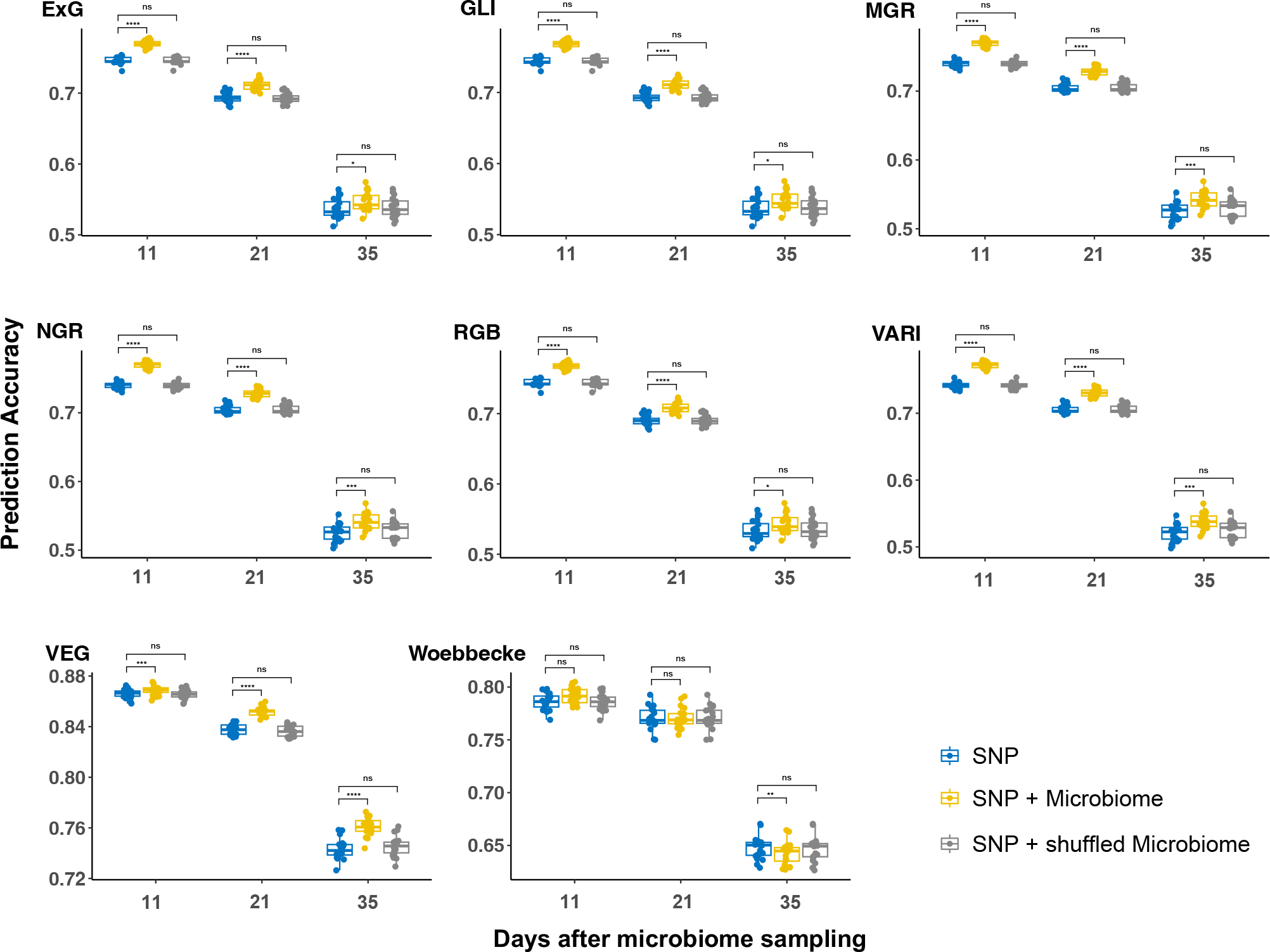
Prediction accuracy of VIs by incorporating microbiome into the prediction model. Genomic prediction results using SNPs only (blue), both SNPs and microbiomes (yellow), and both SNPs and shuffled microbiomes (grey). The statistical significance of the difference in accuracy between the models is indicated by asterisks: ns (not significant), * (0.01 < p-value ≤ 0.05), ** (0.001 < p-value ≤ 0.01), ***(0.0001 μp-value ≤ 0.001), and **** (p-value ≤ 0.0001).

**Figure S4.**
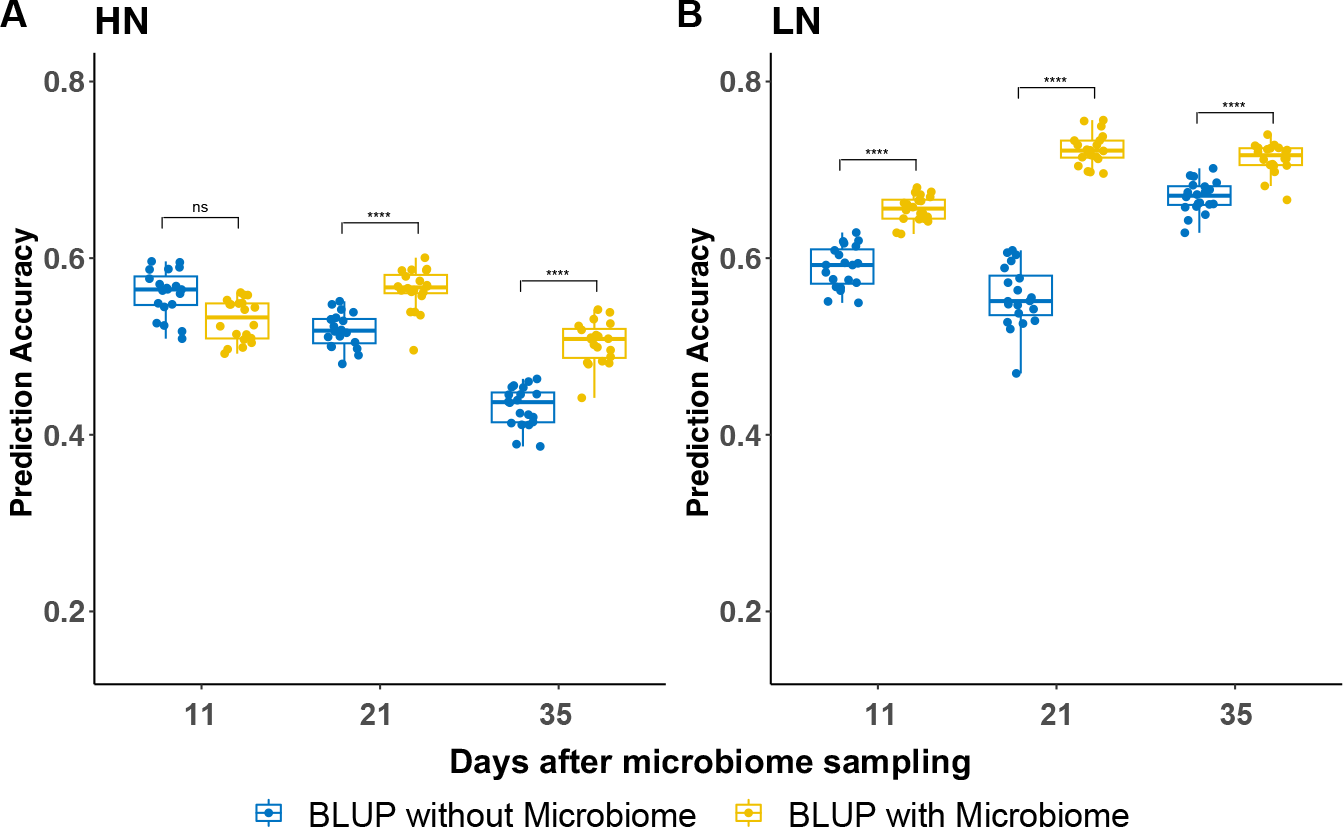
Prediction accuracy of incorporating microbiome under different N conditions. The prediction accuracy of model 1: getting BLUP excluding microbiome effects (blue) and model 2: getting BLUP including microbiome effects (yellow) in high N (**A**) and low N (**B**) conditions. The statistical significance of the difference in accuracy between the models using one tailed t-test is indicated by asterisks: ns (not significant), * (0.01 < p-value ≤ 0.05), ** (0.001 < p-value ≤ 0.01), ***(0.0001 < p-value ≤ 0.001), and **** (p-value ≤ 0.0001).

**Figure S5.**
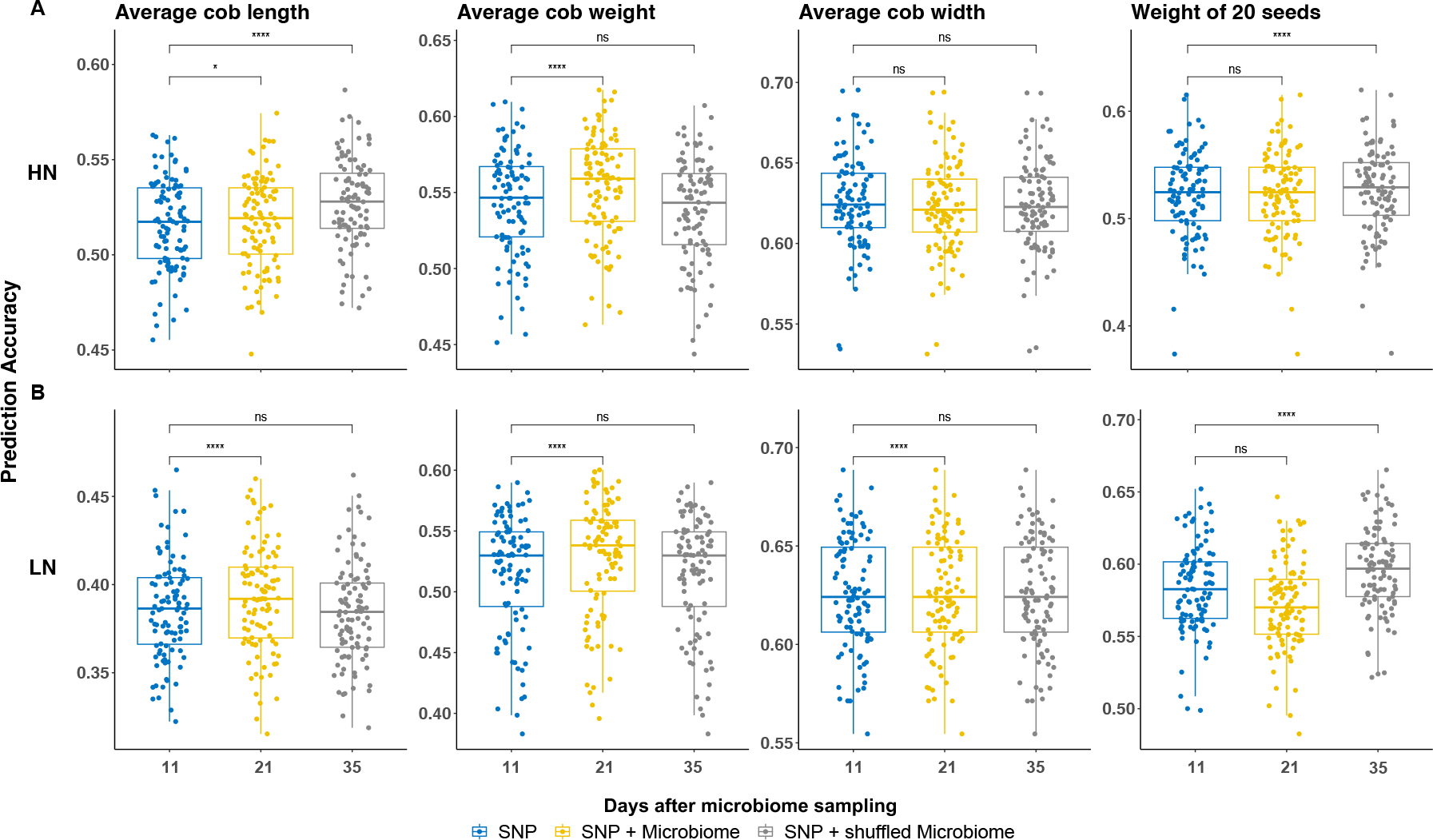
Prediction accuracy for yield-related traits under different N conditions. Genomic prediction results using SNPs only (blue), both SNPs and microbiomes (yellow), and both SNPs and shuffled microbiomes (grey) in the case of high N (**A**) and low N (**B**) conditions. Asterisks indicate the statistical significance of the difference in accuracy between the models: ns (not significant), * (0.01 < p-value ≤ 0.05), ** (0.001 < p-value ≤ 0.01), *** (0.0001 < p-value ≤ 0.001), and **** (p-value ≤ 0.0001).

**Figure S6.**
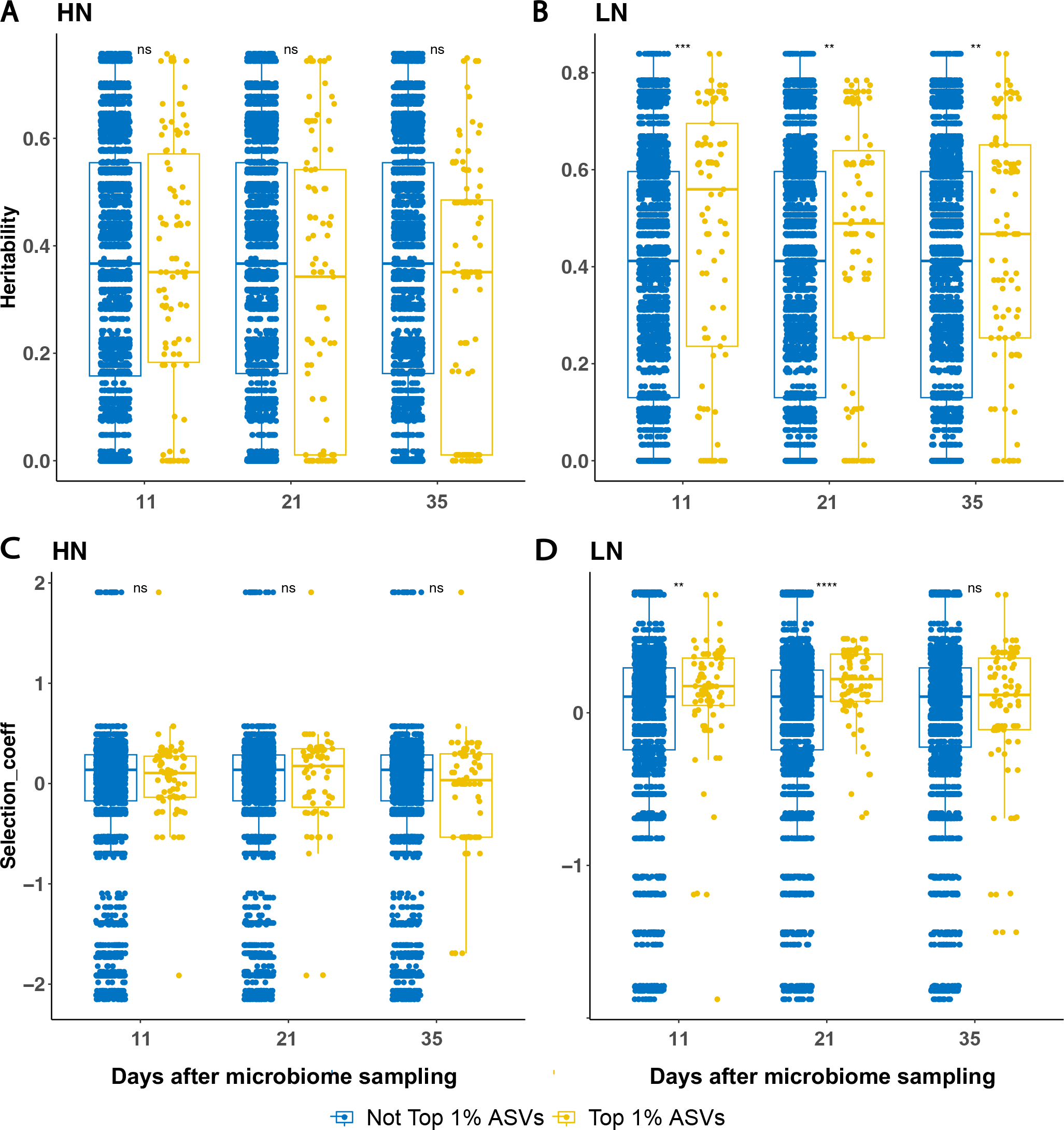
Comparison between top 1% ASVs and the remaining ASVs for heritability and selection coefficient. The Wilcoxon rank sum and signed rank test results between top 1% ASVs (yellow) and the remaining 99% ASVs (blue) for heritability under HN (**A**) and LN (**B**), and selection coefficient under HN (**C**) and LN (**D**), respectively. The statistical significance of the difference of the Wilcoxon rank sum and signed rank test result is indicated by asterisks: ns (not significant), * (0.01 < p-value ≤ 0.05), ** (0.001 < p-value ≤ 0.01), ***(0.0001 < p-value ≤ 0.001), and **** (p-value ≤ 0.0001).

**Figure S7.**
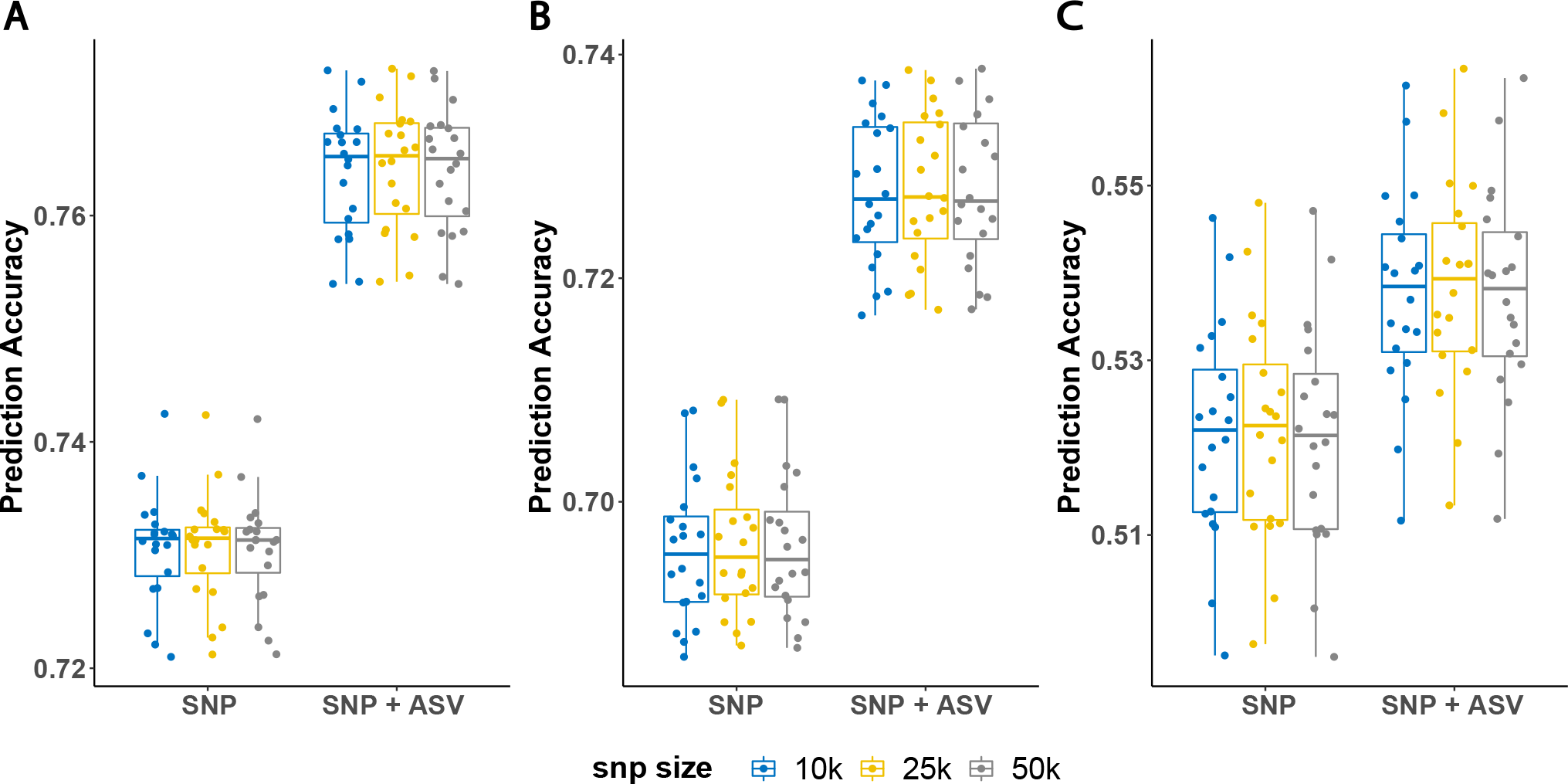
Sensitivity test of the SNP size on prediction accuracy for VARI trait. The Student’s t-test results using randomly selected 10k SNPs (blue), 25k SNP (yellow), and 50k SNP (grey) in 11 (**A**), 21 (**B**) and 35 (**C**) days after microbiome sampling. The test results are not significant with p-values greater than 0.1.

